# Modeling in vitro cell-to-cell spread of hepatitis C viral infection using an agent-based approach

**DOI:** 10.64898/2026.06.05.730411

**Authors:** Zhenzhen Shi, Adam Burns, Evan Cudone, Alexandra Kamm, Karina Durso-Cain, Nicholson Collier, Jonathan Ozik, Susan L. Uprichard, Harel Dahari

## Abstract

Mechanisms that lead to viral chronicity are poorly understood, but cell-to-cell spread has been implicated in the establishment of chronic infections. We previously developed mathematical models to explore the nature of hepatitis C virus (HCV) cell-to-cell spread *in vitro* and quantified the effect of inhibiting individual host factors involved. However, the previous models were not designed to (i) address cell proliferation, (ii) account for differences in cell size, and (iii) did not include possible foci merging. Herein we have developed an agent-based model (ABM) to simulate HCV cell-to-cell spread in vitro by modeling individual cell behaviors. This model recapitulates the natural increase of cell confluence that occurs in vitro accompanied by a concomitant decrease in cell size by allowing for independent proliferation cycles of individual cells within a restricted space. The model fits the experimental foci expansion data well and allows assessment of foci merging while reproducing the irregular HCV foci shape observed in cell culture. Altogether, the new more inclusive model has the potential to help elucidate the dynamics of HCV cell-to-cell spread and provide accurate predictions regarding the efficacy of antiviral drugs.

**Author Summary:** Despite remarkable progress in treatments for hepatitis C virus (HCV), HCV infection remains a global public health burden with over 50 million chronic HCV infections and about 1 million new infections occurring annually. Once infection occurs, HCV can spread in the liver multiple ways. One mechanism is cell-to-cell (CTC) spread where the virus moves directly from one infected cell to an adjacent cell without moving through the extracellular space. Previous mathematical models of HCV CTC spread were not designed to incorporate cell proliferation, cell size, or foci merging. Therefore, to better mimic cells in culture, we developed a novel agent-based model (ABM) that allows the simulated cells to proliferate resulting in an increased number of cells with concomitant decrease in cell/agent size analogous to what happens as cells become tightly packed in culture. This new ABM not only can be used to estimate efficacy values of HCV cell-to-cell spread inhibitors (e.g., when different factors involved in cell-to-cell spread are blocked), but also should enable modeling of HCV CTC spread under a wider variety of cell culture conditions and thus help elucidate the impact of different viral-host dynamics on HCV CTC spread.

## INTRODUCTION

While initial hepatitis C virus (HCV) infection occurs when a virus particle attaches to the outside of a hepatocyte and is endocytosed through the interaction with a specific set of cellular receptors, subsequent infection of naive cells, occurs through two different routes: “cell-free” spread and “cell-to-cell” (CTC) spread. Cell-free spread as described above is characterized by a virus particle egressing from an infected cell and moving freely through the extracellular space before infecting another cell. In contrast, CTC spread is when a virus is transmitted to adjacent cells without ever moving freely through the extracellular space. Many viruses including HCV have been shown to utilize CTC spread as a mode of transmission (1–3). Importantly, because CTC spread protects a virus from antibodies and other extracellular immune factors, it is thought to be involved in the establishment and maintenance of viral persistence and provides an alternative route for the amplification of drug resistant mutants during treatment (4). As such, CTC spread represents a valuable antiviral target. Notably however, current antiviral drug development tends to focus on inhibiting initial entry or viral replication, overlooking CTC spread as a potential antiviral target.

There are many mechanisms by which viruses have been found to spread CTC (3); however, the mechanism and dynamics of HCV CTC spread remain unknown. Several of the factors involved in cell-free HCV entry have been shown to also be involved in HCV CTC spread. For example, blocking of the HCV entry factor Nieman-Pick C1 like 1 (NPC1L1) or Claudin 1 (CLDN1) potently inhibits not only initial cell-free HCV entry, but also subsequent CTC spread (1, 2, 5). Thus, these and other cellular factors involved in both means of HCV spread represent promising antiviral drug targets.

We previously used two mathematical models to explore the kinetics of HCV CTC spread: the *birth* model that assumed that each infected cell in a focus can give rise to another infected cell (as would likely be the case when foci are small) and the *boundary* model where only infected cells at the perimeter of the focus can give rise to another infected cell (as would likely be the case as foci become larger) (6). These efforts quantified the effect of inhibiting individual host factors and reported a hierarchy of efficacies for blocking HCV CTC spread when targeting different host factors involved. While the *birth* and *boundary* models utilized were able to describe the upper and lower limits of the HCV foci growth rates (i.e., cell-to-cell spread) observed, the population-based kinetics of these mathematical models make them unable to account for cell-division, cell size, changes in spread dynamics as foci size changes, or foci merge. Therefore, here we have developed an agent-based model (ABM) to more accurately describe HCV CTC spread on a single-cell basis. Importantly, we have adapted the standard agent-based approach to include agent-division, thus mimicking cell proliferation and the concomitant decrease in cell size that occurs *in vitro* as the monolayer becomes more densely populated.

## MATERIALS & METHODS

### Cells and Virus

Huh7 cells were cultured in Dulbecco’s modified Eagles medium (DMEM) supplemented with 10% fetal bovine serum, 100units/ml penicillin, 100mg/ml streptomycin and 2 mM L-Glutamine. Methods for generation and production of HCV cell culture (HCVcc) stocks from the Japanese fulminant hepatitis (JFH-1) virus clone have been previously described (7).

### HCV spread assay

Experiments and reagents were previously described (6), but in brief, Huh7 cells were plated and infected with 50 focus forming units (FFU) of HCVcc. After 17 hours, the viral inoculum was removed and medium containing 10 μg/mL of the HCV E2 targeted neutralizing antibody AR3A (anti-E2) was added to block cell-free virus spread. As indicated, additional treatments to block HCV CTC spread were added simultaneously with anti-E2 treatment (**Fig. 1**). Treatments included antibodies against claudin-1 (anti-CLDN1), Niemann-Pick C1-like 1 (Anti-NPC1L1), transferrin receptor 1 (anti-TfR1) as well as the inhibitors again TfR1 (ferristatin) and NPC1L1 (ezetimibe). At 72-hour post-inoculation, triplicate wells were fixed with 4% paraformaldehyde and immunostained for HCV E2. The number of HCV-positive cells per focus were counted as a readout of HCV CTC spread. To determine the degree of cell division during the assay period, cell counts from parallel wells were performed at the time of infection and at the time of fixing. Details about the individual treatment and immunohistochemical staining have been previously described (6).

**FIG 1.**
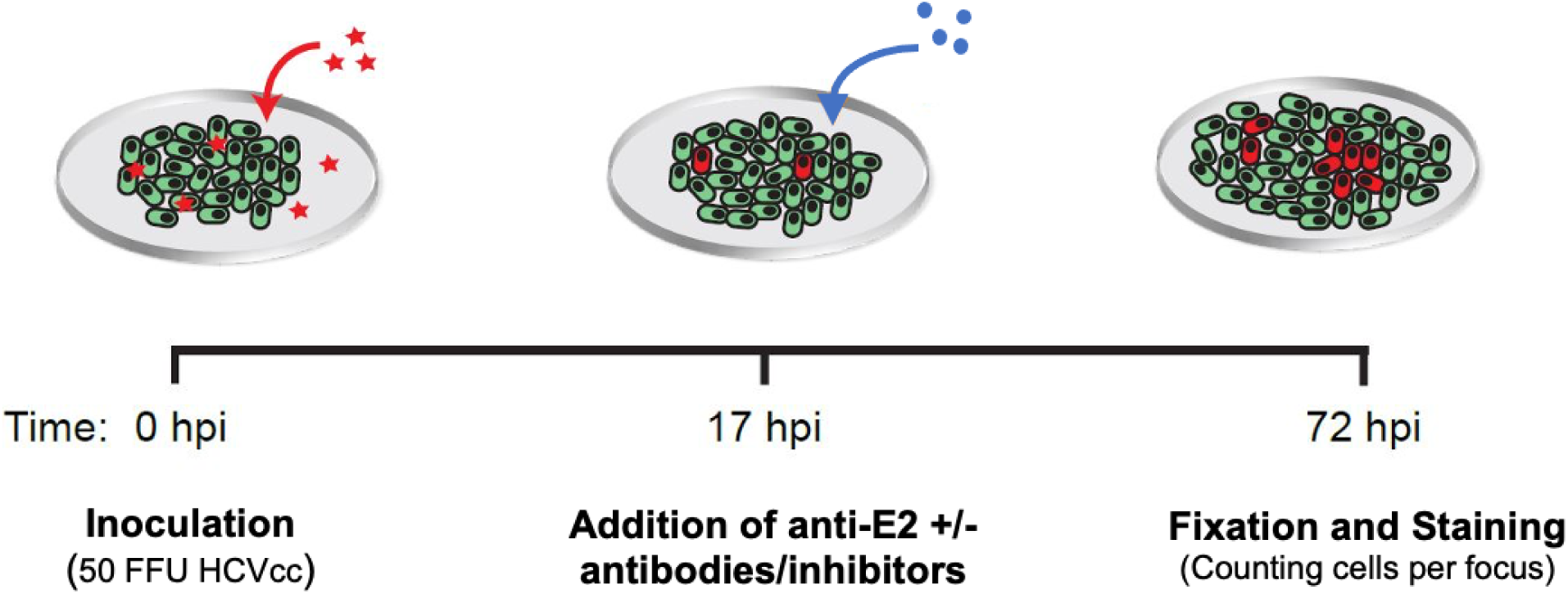
HCV spread assay. Huh7 cells were plated and infected with 50 focus forming units (FFU) of HCVcc. After 17 hours, medium containing 10 μg/mL of the HCV E2 targeted neutralizing antibody AR3A (anti-E2) was added to block cell-free virus spread. Additional treatments including anti-CLDN1, anti-NPC1L1, anti-TfR1, ferristatin, and ezetimibe were simultaneously added to block CTC virus spread. At 72-hour post-inoculation (hpi), the number of HCV-positive cells per focus were counted as a readout of HCV CTC spread. Uninfected cells are green. Infected cells are red. Red stars: HCV virus. Blue solid circles: anti-E2 and additional treatments including anti-CLDN1, anti-NPC1L1, anti-TfR1, ferristatin, and ezetimibe.

### Modeling cell division

Because the experimental data indicates that cell division occurred during the HCV CTC spread assays (6) and viral spread involves movement between cells, we chose to develop an ABM in AnyLogic (version 8.9.4) which employs a 2-dimensional grid that mimics a monolayer of cells in an *in vitro* cell culture dish (**Fig. 2A**). This ABM represents roughly 1/5 of the total cell population from a single well of the HCV spread assay described above (2000 cells). Agents represent units of space in the model, which tile a 2D grid representing the growth surface. Each of these agent tiles is occupied by 0, 1, 2, 3, or 4 cells, with the number increasing as the cells divide within the constant tile space (**Fig. 2B**). Therefore, this models the compaction of the cells observed in culture without changing the size of the agents. Each of up to 4 individual cells in a single agent can be infected independently and infect the other cells within the agent and other cells in neighboring agents independently. Experimentally the *in vitro* cell cultures experienced a roughly 5-fold increase in population during the 72-hour experiment, thus we have parameterized the ABM to match this using two proliferation mechanisms: proliferation into empty space and compaction.

**FIG 2.**
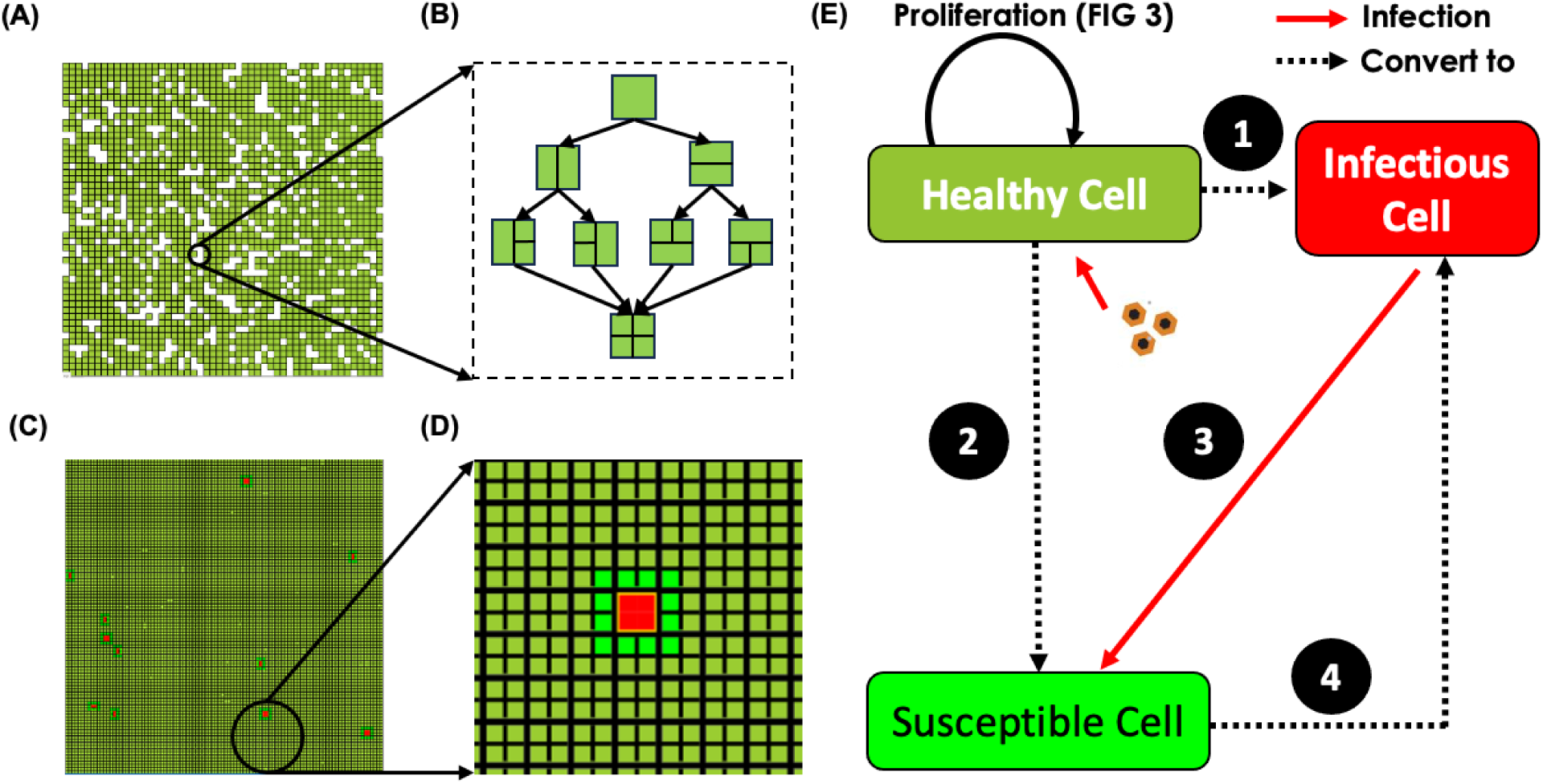
Modeling cell proliferation and cell-to-cell infection spread. (A) Initial grid of 50×50 cell blocks (S0 and S1A) at time 0.0 hours. Dark green cells represent uninfected susceptible cells and white cells represent empty space. (B) A green square in an ABM represents an Huh7 cell plated in a monolayer. Each cell agent represented by a square can divide into 2, 3, or 4 separate cells without changing the size of the original cell to account for the compaction of cells observed in culture. (C) Final grid of 100×100 possible cell blocks with initially infected cells (red cells) without spread at 72 hours post-inoculation. (D) The red represents infected cells, the lime green represents adjacent susceptible cells, and the dark green represent the non-adjacent non-susceptible cells. The yellow outline shows the first cell infected in the foci, i.e., the foci origin. In the simulations, infected cells were not allowed to proliferate and thus remain undivided, as can be seen by the infection origin. (E) Infection spreads through cell states. 1: First, cells are infected by cell-free viral particles added to the medium. These infection events occur randomly at a rate of 𝜆(𝑡) = 𝜆_0_(𝑒^―𝛼𝑡^) with an initial infection rate 𝜆_0_ = 6.59 foci per hour and infectivity loss 𝛼 = 0.065 per hour; 2: Cells become susceptible to infection by being immediately adjacent to cells in the infectious state; 3: Infected cells are capable of transferring infection to healthy neighboring cells at an infection spread rate (i.e. ISR in **Table 2**). 4: Susceptible cells then become infectious and are capable of transferring infection to healthy neighboring cells (step 2).

### Modeling proliferation of cells into empty space

To match the 5-fold increase in cell number that occurred experimentally, the ABM is initiated by seeding 80% of the agent tiles with a single cell (**Fig. 2A**, 2000 green cells). This leaves room for *in silico* cells to proliferate into 20% empty space (**Fig. 2A**, 500 white cells). Since the generation of each new cell in an empty space is assumed to be independent of one another an exponential distribution was used in the form of a Poisson point process. This model proliferation mechanism is illustrated in **Fig. 3** by the transition from states S0 to S1B (**Table 1**).

**FIG 3.**
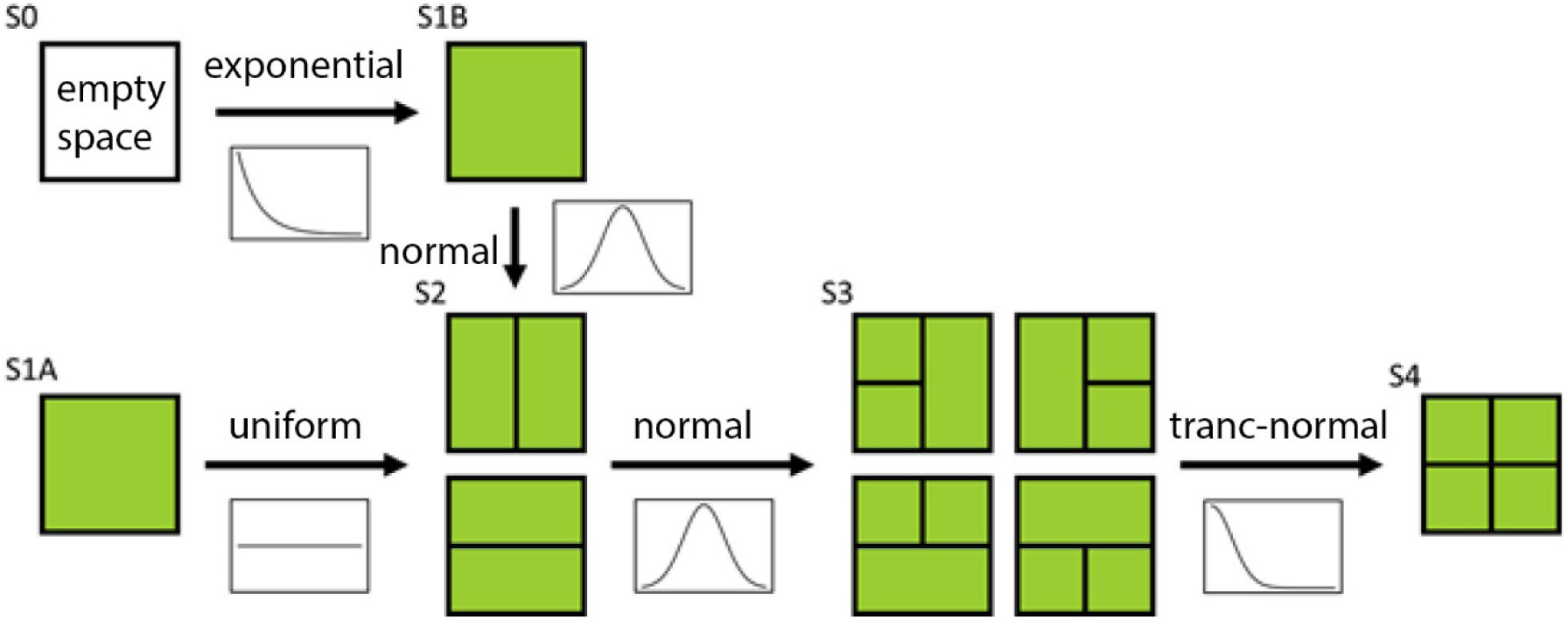
Depiction of the *in-silico* cells and the time distributions for each proliferation state. Green squares/rectangles represent cells. S0, S1A, S1B, S2 and S3 indicate the different cell states. Arrows indicate a cell division and the time distribution for each division is indicated below the arrows. **S0** represents empty (white) space in the grid. **S1A** represents susceptible cells (green) at initiation time of experiment. **S0 to S1B:** represents the generation of new *in-silico* cells in the empty space of the grid (Fig. 2A), without changing cell size via exponential time distribution. **S1B to S2:** represents the *in-silico* cell proliferation of S1B into 2 smaller daughter cells via a normal-time distribution. **S1A to S2:** represents the *in-silico* cell proliferation of S1A into 2 smaller daughter cells via uniform time distribution. **S2 to S3:** represents the *in-silico* cell proliferation of the first S2 cell into 2daughter cells via a normal time distribution resulting in an agent with 3 cells. **S3 to S4:** represents the second S2 cell into 2 daughter cells via truncated-normal-time distribution resulting in an agent with 4 cells. Detailed explanation on assumed time distributions and ranges are provided in Methods and **Table 1**.

**TABLE 1.** Model characteristics.

| Characteristic | Cell state<br>(Fig. 3) | Default value<br>[unit] | Distribution |
| --- | --- | --- | --- |
| Cell proliferation to empty space<br>(min, lambda) | S0 to S1B | (0, 20)<br>[hr] | Exponential |
| First division of initial cell<br>(max, min) | S1A to S2 | (24, 0)<br>[hr] | Uniform |
| First division of non-initial cell<br>(mean, SD) | S1B to S2 | (24, 3)<br>[hr] | Normal |
| Second Division<br>(mean, SD) | S2 to S3 | (24, 3)<br>[hr] | Normal |
| Second Division #2<br>(mean, SD) | S3 to S4 | (0, 3)<br>[hr] | Truncated Normal |
| Infection spread rate | NA | 10 | NA |

Note: SD - standard deviation; NA - not applicable

### Modeling compaction of proliferated cells

To account for cell division after confluence, we incorporated another *in silico* proliferation mechanism to allow cells to subdivide 2 more times while continuing to occupy the parent cell’s original space (**Fig. 2B**). During the *in-silico* cell division process, each cell maintains an individual cellular clock with time of division determined by a set distribution (**Table 1**). Following this predetermined distribution, cells at different states have explicit proliferation rules. Cells in state S1A or S1B are able to divide resulting in 2 daughter cells (**Fig. 3**, state S2). The division resulting from S1A occurs via a uniform time distribution between 0 and 24 hours since the age of cells at the time of initiation is not known. However, the division resulting from S1B occurs via a normal time distribution with a mean of 24 hours because the age of each cell is known. Because the S2 cells may not divide at the same time, we represent the final cell division occurring first in one of the S2 daughter cells (S3) and then in the second S2 cells leading to the final S4 state. The division resulting from cells in state S2 to cells in state S3 occurs via a normal time distribution with a mean of 24 hours. However, due to a modeling limitation, once one of those two cells in S2 divides, it resets the clock for all the cells in its parent cell i.e., the daughter cells that occupy the space of the original cell size. To account for that, a truncated-normal-time distribution with an average rate of 3 hours was chosen for the transition from S3 to S4.

### Modeling HCV cell-to-cell spread

Cell infection by HCV in the simulation is tracked through three infection states: susceptible but not adjacent to infected cells (dark green), susceptible and adjacent to infected cells (fluorescent green), and infected (red) (**Fig. 2C** and **2D**). All cells are initiated in the susceptible state. *In silico HCV* infection of susceptible cells occurs by two mechanisms: 1) the initial cell-free virus infection that occurs before the anti-E2 introduction at 17 hours post-inoculation and 2) CTC virus infection after anti-E2 addition (**Fig. 2E**). Initial cell-free viral infections happen stochastically, infecting random cells individually. Any individual cell, regardless of proliferation state, can be the target of initial cell-free virus. As previously shown in Graw et al. (6) the initial cells-free infections occur at a rate of 𝜆(𝑡) = 6.59(𝑒^―0.65𝑡^) and these initially infected cells are indicated in the simulation by a yellow outline (**Fig. 2D**). The second mechanism of infection in the model, CTC infection, occurs stochastically between an infected cells and one of its adjacent Moore neighbors. The adjacent Moore neighbors become susceptible to HCV and are infected at an infection spread rate per simulation step, where 1 simulation step represents 1 hr in the experiment. Since a single cell agent can have up to 4 individual cells within it (**Fig. 3**, state S4), the state of each daughter cell within a single cell agent is explicitly evaluated in the simulation. Once the cell gets infected, its Moore neighbors will become susceptible to HCV infection (**Fig. 4**, lime green). The foci size (i.e., the number of infected cells within each HCV-positive foci of cells) and the number of foci in the grid/monolayer are calculated at the end of the simulation. Infected cells divide slower than uninfected cells (8), however, for simplicity we assume in our model that once a cell is infected it can no longer proliferate. We also assume that no cell death occurs in our model based on the lack of any observed cell death during the 72h assay.

**FIG 4.**
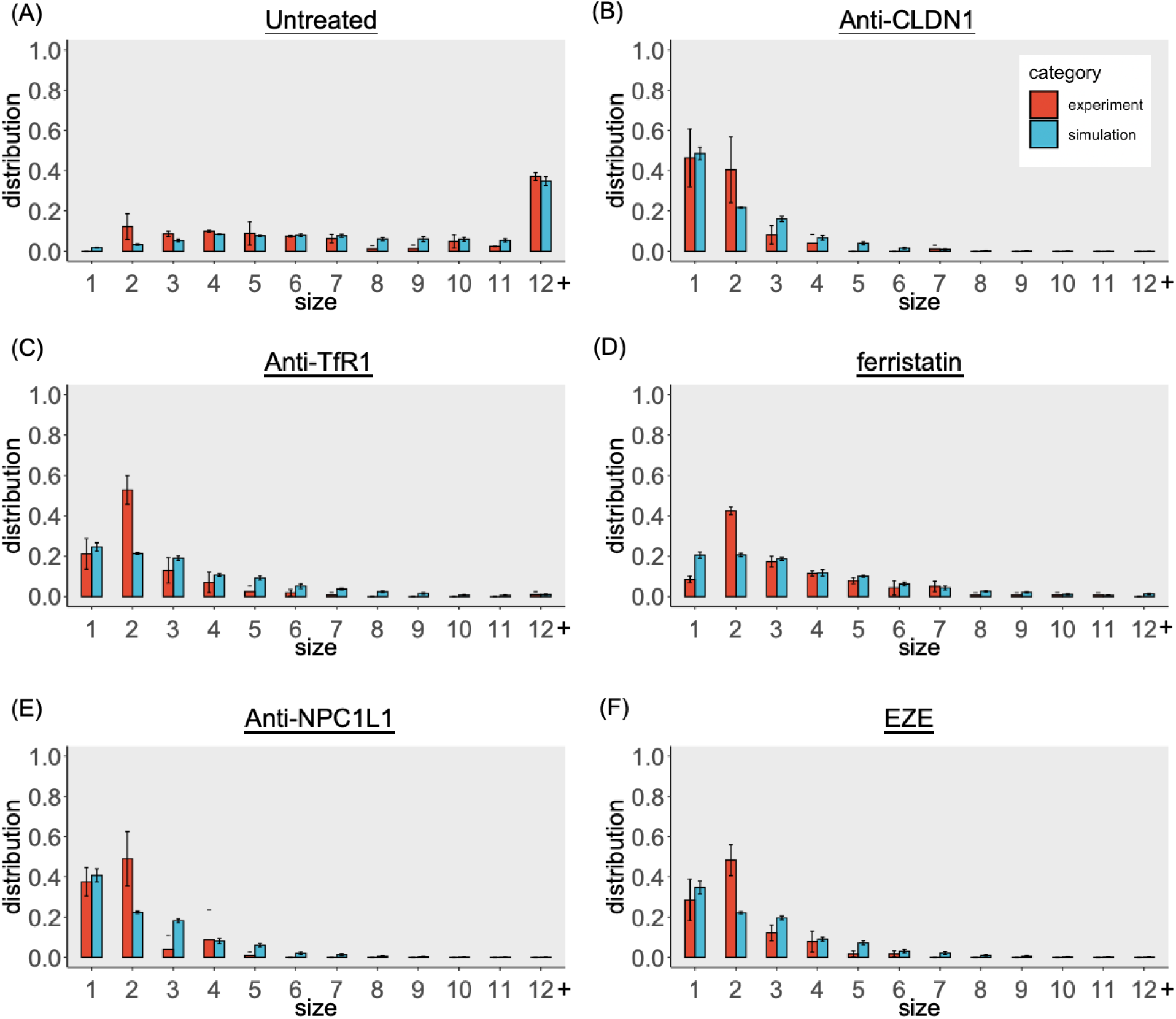
ABM calibration with experimental data. The distribution of focus sizes in the CTC spread assay at 72 h post-inoculation without **(A)** and with inhibitors **(B-F)** that target cellular HCV entry factors CLDN1 (anti-CLDN1), NPC1L1 [anti-NPC1L1; ezetimibe (EZE)], or TfR1 (anti-TfR1; ferristatin). Foci comprising 12 or more cells were combined into the ‘12+’ cells category. The experimental data is an average of 2 wells for untreated condition and an average of 3 wells for inhibitors conditions (red bars). The simulation results (blue bars) are an average of foci distribution calculated using the best GA estimates (Table 2 and Table 3). Error bars indicated mean ± SD.

**TABLE 2.** ABM best fit parameter estimates for HCV CTC spread. Genetic algorithm (GA) was used to fit the untreated (i.e., uninhibited) HCV CTC spread data from triplicate wells (**Fig. 4A**). The objective function of GA is to minimize the difference between experimental data and simulated results. The eleven parameter combinations that fit all targets in GA are shown. The range of fits (mean ± SD) for each foci size distribution was calculated using all parameter combinations with 100 random seeds (0–99). ISR - infection spread rate; CPR - cell proliferation rate; FDTIM - maximum time for the first division of initial cells; FDTNM - mean time for the first division of non-initial cell; SDTM - mean time for the second division.

| Parameter combination # | ISR [/h] | CPR [h] | FDTIM [h] | FDTNM [h] | SDTM [h] |
| --- | --- | --- | --- | --- | --- |
| 1 | 10.73 | 25.13 | 27.06 | 24.03 | 28.85 |
| 2 | 10.73 | 25.13 | 27.06 | 24.52 | 29.64 |
| 3 | 10.73 | 25.13 | 27.06 | 31.52 | 28.85 |
| 4 | 10.73 | 25.13 | 27.06 | 27.83 | 28.85 |
| 5 | 10.73 | 25.13 | 27.06 | 24.52 | 30.03 |
| 6 | 10.86 | 12.96 | 34.31 | 23.20 | 28.04 |
| 7 | 10.73 | 13.72 | 12.67 | 16.15 | 34.52 |
| 8 | 10.21 | 27.91 | 29.74 | 23.12 | 28.85 |
| 9 | 10.86 | 14.12 | 34.31 | 35.88 | 28.04 |
| 10 | 10.78 | 12.53 | 28.77 | 33.41 | 16.84 |
| 11 | 10.08 | 16.94 | 22.86 | 18.16 | 28.81 |

**TABLE 3.**
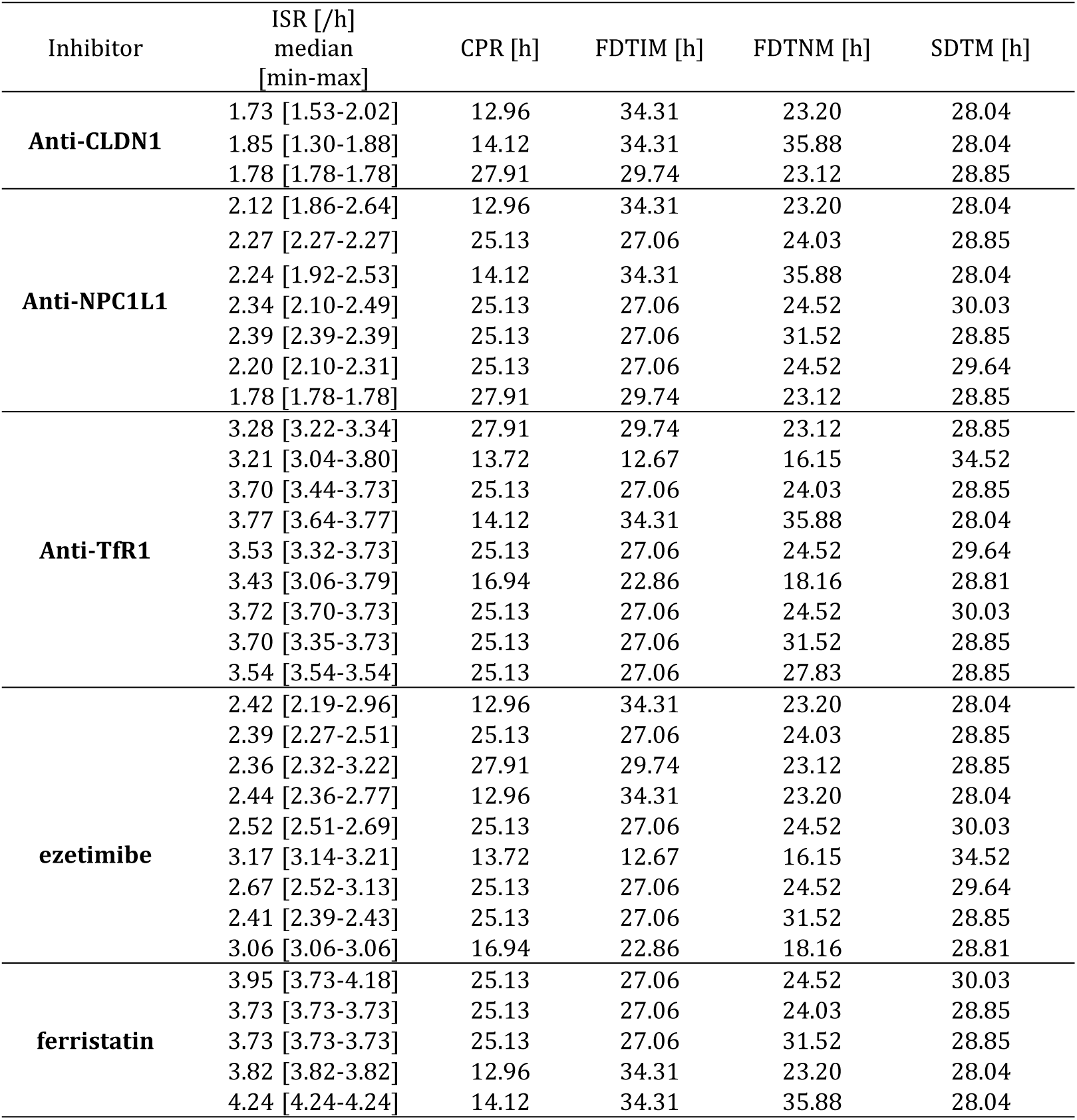
ABM best fit parameter estimates for HCV CTC spread under different inhibitor treatments. . Genetic algorithm (GA) was used to fit the HCV CTC spread data under different inhibitor treatments (**Fig. 4B-F**). Parameters CPR, FDTIM, FDTNM, SDTM were fixed using the GA fit estimated values determine for the untreated condition (Table 2). ISR was fitted to the inhibitor data from triplicate wells. The top 1000 parameter combinations with the smallest Mean Squared Errors for each of the five inhibitors were selected and are summarized. Repetitive parameter combinations during the sampling were removed.

### Parameter estimation

Model parameter fitting was done using a Genetic Algorithm (GA) (9) with the EMEWS framework (10) on Bebop, a high-performance computing (HPC) cluster administered by the Argonne National Laboratory Computing Resource Center. The fitting was performed in two steps. In the first step, a range of five key parameters (**Table 2**) was used to find optimal model parameter combinations for fitting foci size distribution obtained from the anti-E2 treatment. We sampled five parameters from predefined ranges (**Table S1**), where each parameter combination was run using 100 different random seeds (0–99). The objective of the GA was to find the lowest mean squared error (MSE) with the empirical calibration targets of foci size distribution across the model parameters, where each parameter’s distribution of foci sizes was calculated as the mean across 100 replicates. Eleven parameter sets that produced outputs within the bounds for the empirical foci size distribution were identified and are listed in **Table 2**. In the second step, we fit the data from the remaining five individual treatments (anti-CLDN1, anti-TfR1, ferristatin, anti-NPC1L1, and ezetimibe (EZE)]) using the GA in eleven separate experiments for each treatment. In each experiment, we sampled the infection spread rates (i.e., ISR) parameter from a predetermined range of 1.0 – 20.0 and set the remaining four parameters to corresponding values from one of the eleven best anti-E2 parameter sets. As in the first step, each experiment was run with 100 random seeds, and the objective of the GA was to find the least MSE across the seeds. The best parameter combinations across the eleven experiments for each treatment are shown in **Table 3**.

The GA was implemented using the DEAP (11) evolutionary computation Python framework (specifically (12): Chapter 7) and integrated into an EMEWS HPC workflow using EMEWS queues for Python (EQ/Py) (13, 14). The use of HPC resources enable the concurrent evaluation of large numbers of design points, reducing the time to solution. During each iteration of the GA, the best points from the currently evaluated population were selected using a tournament selection method to create a new population. Each of these points was then “mated” with another according to a crossover probability and, finally, each of the resulting points was mutated according to a mutation probability. At each GA algorithm iteration, the new population was evaluated in parallel and the evaluation results were gathered. In the first step, the GA population size was set to 1300, the mutation probability to 0.2, the crossover probability to 0.5, and the number of iterations to 20. The runtime for a typical first step run was 7 hours using full concurrency on 10 nodes (with 36 cores per node), or about 2520 core hours. In the second step, the GA population was set to 100, the mutation probability to 0.5, the crossover probability to 0.5, and the number of iterations to 10. The runtime for a typical second step run was 1.25 hours using full concurrency on 3 nodes (with 36 cores per node), or about 135 core hours.

## RESULTS

### The ABM is able to reproduce the number and size of foci under baseline conditions and during inhibition of CTC spread

To determine whether the new ABM that simulates individual cell division and virus CTC transmission can accurately reproduce the observed foci size distribution (i.e., HCV cell-to-cell spread), we varied five key parameters to fit the ABM to the experimental data under the baseline CTC assay conditions where only anti-E2 alone is included to block HCV cell-free spread (see Materials and Methods). The best model fits which minimized the difference between simulated results and experimental data were selected (**Table 2**). The ABM reproduced the observed *in vitro* data well (p= 0.5361; Kolmogorov-Smirnov test) indicating foci size distribution in the ABM is similar to that obtained experimentally under baseline conditions when CTC occurs uninhibited (**Fig. 4A**). Interestingly, unlike our prior models (6), the new ABM also reproduces the irregular foci shapes reminiscent of what is observed in cell culture (**Fig. 5A-E)**.

**FIG 5.**
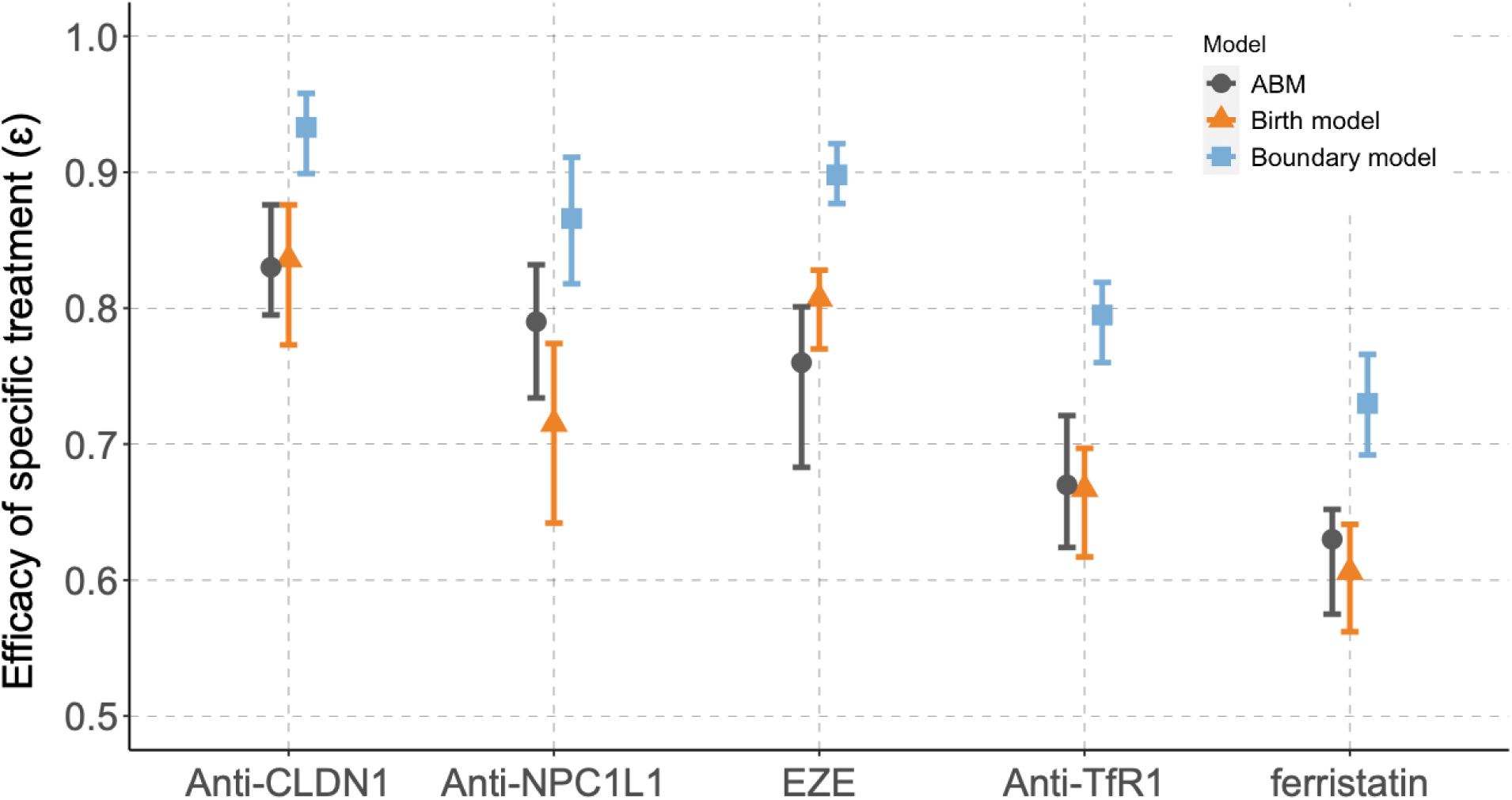
Agent-based vs mathematical modeling efficacy estimation of HCV cell to cell spread antibodies and inhibitors. Graw et al. (6) *birth* (orange) and *boundary* (blue) stochastic models calibration results were plotted against the current ABM results (black). The triangles and squares indicate the estimated effectiveness of specific treatments (ɛ) from the *birth*/*boundary* models and the ABM, respectively. The error bars indicate 10% and 90% percentiles for the *birth*/*boundary* models and the minimal and maximal for the ABM as shown in **Table 4**.

**TABLE 4.** Comparison of treatment efficacy estimates between models. . The efficacy (ɛ) of each HCV CTC spread inhibitor estimated using the new ABM and our previously reported *birth* and *boundary* models is shown. The models were fit to the average of triplicate experimental wells. Numbers in parentheses for the birth and boundary models are the 10% and 90% percentiles over 200 bootstrap replicates of the data, (Table 3 in Graw et al.(6)). Numbers in parentheses for the ABM are the minimal and maximal derived from best fits in Table 2 and Table 3.

| Factor | birth model<br>$\varepsilon$ (%) | ABM<br>$\varepsilon$ (%) | boundary model<br>$\varepsilon$ (%) |
| --- | --- | --- | --- |
| Anti-CLDN1 | 83.6 (77.3, 87.6) | 83.4 (80.0, 88.0) | 93.3 (89.9, 95.8) |
| Anti-NPC1L1 | 71.5 (64.2, 77.4) | 79.4 (73.8, 83.6) | 86.6 (81.8, 91.1) |
| Anti-TfR1 | 66.7 (61.7, 69.7) | 66.9 (62.3, 72.0) | 79.5 (76.0, 81.9) |
| ezetimibe | 80.7 (77.0, 82.8) | 75.7 (68.0, 79.8) | 89.8 (87.7, 92.1) |
| Ferristatin | 60.6 (56.2, 64.1) | 63.4 (57.9, 65.6) | 73.0 (69.2, 76.6) |

To fit the model to the CTC spread data in the presence of the inhibitors, we only varied virus spread rate since the inhibitors were used at concentrations that do not alter cell division (**Table 3**). Again, the ABM fit the experimental data (p > 0.5; Kolmogorov-Smirnov test) (**Fig. 4B** – **4F**, blue vs red bars, respectively). The estimated ISR in the presence of the inhibitors are much smaller (1.7 - 4.2) compared to the estimated ISR in the absence of the inhibitors (10.1 – 10.9).

### Modeling HCV spread inhibition to quantify efficacy of cell-to-cell spread inhibition

To quantify the effect of antibodies and inhibitors on CTC spread, we defined a parameter ε to measure the degree to which focus expansion was reduced under different treatment conditions. Consistent with the experimental data and similar to our previous findings using mathematical modeling approaches (6), the same hierarchies of estimated focus growth reduction were found using the ABM when comparing treatment with the neutralizing antibodies (e.g., CLDN1 > NPC1L1 > TfR1) and the small-molecule inhibitors (e.g., EZE > ferristatin) (**Fig. 6** and **Table 4**). Specifically, the ABM results demonstrated that blocking CLDN1 with antibodies resulted in the largest reduction of foci expansion (on average 83% reduction) and small-molecule inhibitor ferristatin exhibited the smallest reduction of 63% on average. As might be expected when assessing relatively small foci, the ABM tended to provide efficacy estimates closer to the *birth* model (6) that assumed each infected cell in a focus can give rise to another infected cell.

**FIG 6.**
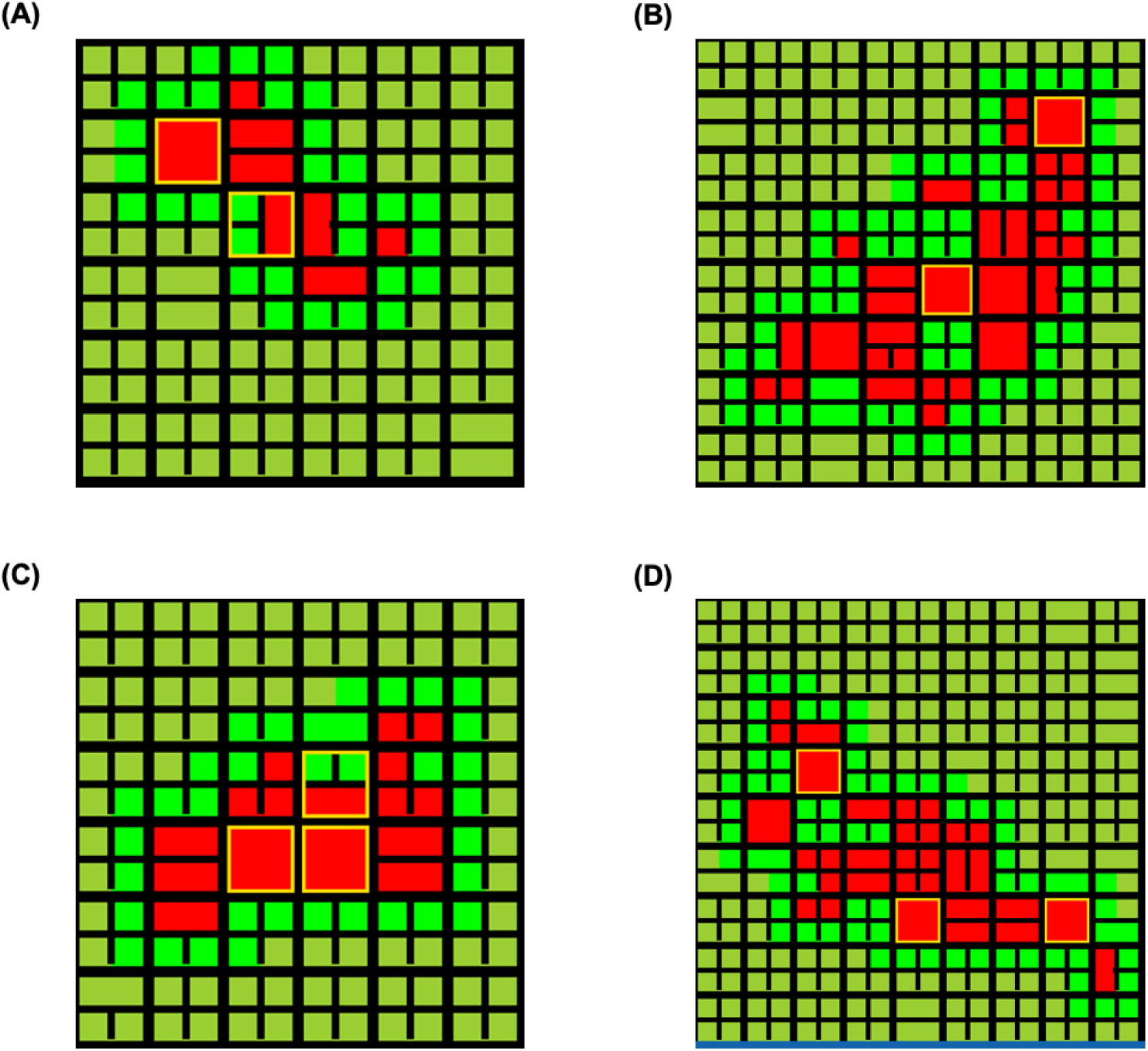
Images of *in-silico* merging foci. (A-B) Merged foci formed from two individual foci (initially infected agent highlighted in yellow) that were initially closely situated (A) or two initially distantly situated (B). **(C-D)** Merged foci formed from three individual foci initially closely situated (C) or initially distantly situated (D). In some cases, a cell had divided before the infection.

### The ABM predicts only one foci merging event in less than half of the cell-to-cell spread simulations

While there is no way to know if any experimental foci observed resulted from merging of initially independent foci, the ABM has the ability to identify whether foci have merged during the simulations by counting the number of initially infected cells (highlighted in yellow) in each focus. Concentrating on the baseline conditions where anti-E2 blocks cell-free spread, but no inhibitors are included, we ran the ABM using the random parameter combination No. 7 (**Table 2)** with 100 different random seeds (0–99) of average of 11 infected cells (which is equivalent to the 50 cells when scaled up to *in vitro* setting). In 100 simulation runs, a single merging foci event, consisting of two individual foci, was identified 43% of the time (43/100). These merged foci were formed by the growth of two individual foci which were initially infected individually (**Fig 7A and 7B**). In 2% of the simulations (2/100) three initially infected cells were found in a single merging foci (**Fig. 7C** and **7D**). Consistent with the reduced foci growth observed in the conditions that include the various inhibitors, foci merging events were rare in the inhibitor simulations. For example, in 100 anti-CLDN1 simulation runs, a foci merging event occurred only 20% (20/100) of the time and no merged foci consisting of three individual foci were predicted.

**FIG 7.**
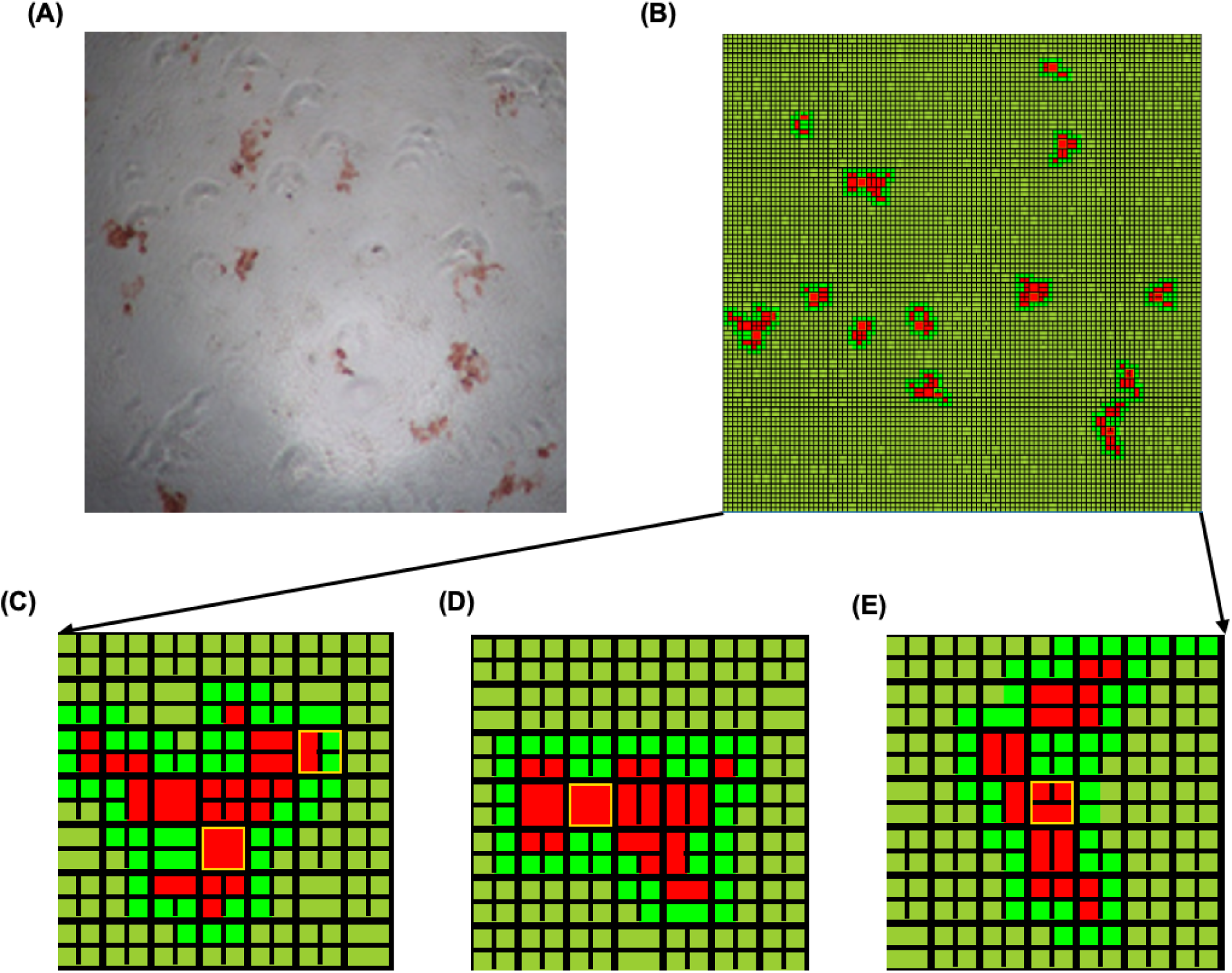
Experimental HCV foci formation and agent-based model simulations. **(A)** Image of experimental HCV cell-to-cell spread assay data. **(B)** Image of foci formed after 72 h post-inoculation *in silico* using the parameter combination **in Table 2** (ISR: 10.86; CPR: 12.96; FDTIM: 34.31; FDTNM: 23.20; SDTM: 28.04) and a random seed of 3. **(C-E)** Image of distinct foci formed *in silico*. The red represents infected cells, the lime green represents adjacent susceptible cells, and the dark green represents remotely susceptible cells as explained in detail in Methods (FIG 2). The yellow outline shows the first cell infected in the foci, i.e., the foci origin. In the simulations, infected cells were not allowed to proliferate and thus remain undivided, as can be seen by the infection origin and the partially divided cells surrounding it (C).

## DISCUSSION

In the present study, we developed an agent-based model to simulate *in vitro* HCV CTC spread. By integrating experimentally measured cell division parameters, the model captures the natural increase in cell confluence alongside a decrease in cell size and allows the model to assess foci merging. Importantly, the new ABM aligns closely with the experimental data and is therefore able to estimate the efficacy of HCV cell-to-cell inhibitors.

Our previous *birth* and *boundary* mathematical models (6) where thought to describe the upper and lower limits of HCV foci growth rates as they were designed such that foci growth could be affected either by all the infected cells in a focus (*birth* model) or only the infected cells at the periphery of the focus (*boundary* model). In contrast, the current ABM was built upon individual-cell simulation that assesses whether an infected cell has an uninfected neighbor, thus allowing for CTC spread to occur in response to foci size and shapes based on stochastic parameters such as infection spread rate and the state of cell division. Overall, using the ABM to evaluate the effect of various inhibitors targeting different host cell factors on cell-to-cell spread, generated the same hierarchy of efficacies as the previous models. However, the ABM provided a fit to the foci size distribution across different treatment conditions which was closer to foci distributions generated by the *birth* model (**Fig. 5**), likely because in the standard 3-day foci assay being assessed, HCV foci remain relatively small, especially under treatment, such that most infected cells are still adjacent to uninfected cells making the *birth* model most relevant. The results may indicate that the *birth* model can be used as a simple modeling approach for short duration experiments.

Our ABM describes different stages of cell division and mimics viral spread through modeling individual cell-to-cell contacts, with the ultimate goal of differentiating the effect of viral spread and cell division on foci growth. However, because infected cells have been observed to divide slower than uninfected cells (8), in this initial study we simplified the modeling by not allowing cells to divide once they are infected. Having removed this direct impact of cell division on foci growth, we performed simulations adjusting uninfected cell division to determine if there were any effects of uninfected cell division on foci growth (**Fig. S1**). While increasing cell-to-cell infection spread rate (i.e., ISR), resulted in increased foci size at 72h post-inoculation compared to the default setting as expected (**Fig S1A vs B**), foci size did not change significantly when different cell division rates were adjusted to make cell division faster (**Fig S1A vs. C-F**). Determining how much infected cell division contributes to foci growth in our *in vitro* cell culture system would be useful when experimentally quantifying CTC spread based on foci size, hence future work will include experimentally determining the rate of division of infected and non-infected cells and adding division of the infected cells into the model.

Another concern when experimentally assessing viral spread based on foci size is the question of foci merging. While sometimes foci shape might suggest the merging of 2 foci and a reduction in foci number over time is suggestive of foci merging, it is not feasible to know if a specific focus is the result foci merging. An advantage of the ABM is that the foci-initiating infected cells are known, allowing foci merging to be followed (**Fig. 7**). Consistent with our effort to avoid foci merging in our experiments, very little foci merging was predicted by the model (i.e., only one merging event in less than 50% of simulations). However, adding in division of infected cells to the simulations as planned is expected to result in an increased rate of foci growth, which in theory will affect the kinetics of foci merging. In the absence of division of infected cells, viral cell-to-cell spread was the major driving force for foci merging as expected. However, as cell division became faster, two individual foci could fail to merge under the same cell-to-cell infection spread rate (**Fig. S2**) presumably because the number of uninfected cells between the initial foci increased creating a larger cell barrier between foci.

A curious observation made in the model simulations is that the foci shape observed in the model simulations are very irregular and similar to the irregular foci shapes observed *in vitro* **(Fig. 5)**. We initially thought that perhaps the simultaneous virus spread and cell division was the primary reason for the irregular shapes, but both *in vitro* (**Fig S3A**) (7, 15) and *in silico* experiments (**Fig. S3B**) under conditions of no cell division still produced irregular shaped foci. Hence, the reason for the unusual foci shape still needs to be determined and could be addressed with future models (e.g., testing different agent shapes/grid patterns).

With the experimental evidence documenting how variable the stochastic nature of viral infection can be (16, 17), it is clear that an ABM approach is better able to simulate infection dynamics, particularly viral spread. We previously published an HCV spread ABM that included both cell-free and CTC spread with the goal of determining the extent to which these two different transmission mechanisms contributed to HCV spread (18). While the prior ABM included a small degree of initial cell division, it was not designed to accurately recapitulate the degree of cell division observed experimentally, nor was it possible to change the rate of cell division. To better understand HCV CTC spread, here we developed an ABM to mimic our *in vitro* HCV CTC spread assay which is designed to eliminate HCV cell-free spread so that HCV CTC spread can be monitored in isolation. This focus allowed us to incorporate a more accurate representation of the cell culture monolayer (e.g., cell division and changes in cell size). Therefore, this new ABM can not only estimate the efficacy of HCV CTC spread inhibitors but also enables modeling of HCV CTC spread across a broader range of cell culture conditions, thereby helping to elucidate how different viral–host dynamics influence HCV CTC spread. In the future, we envision creating a hybrid model combining the most relevant features of these different ABMs with a detailed mathematical representation of intracellular dynamics to describe HCV infection kinetics. This type of approach would be useful for dissecting the dynamics of HCV spread as well as estimating the effectiveness of antiviral drugs.

## ACKNOWLEDGEMENT

We thank Frederik Graw for insightful discussions. This research was completed with resources provided by the Laboratory Computing Resource Center at Argonne National Laboratory.

## Supplementary Material

**TABLE S1.** Parameter range used for Genetic Algorithm (GA)

| Parameter | Symbol<br>[unit] | Range tested<br>for GA step 1 | Range tested for<br>GA step 2 |
| --- | --- | --- | --- |
| Infection spread rate | ISR [/h] | 1-20 | 1-20 |
| Cell proliferation rate | CPR [h] | 10-30 | NA |
| Maximum time for the first division of<br>initial cells | FDTIM [h] | 12-36 | NA |
| Mean time for the first division of<br>non-initial cell | FDTNM [h] | 12-36 | NA |
| Mean time for the second division | SDTM [h] | 12-36 | NA |

**FIG S1.**
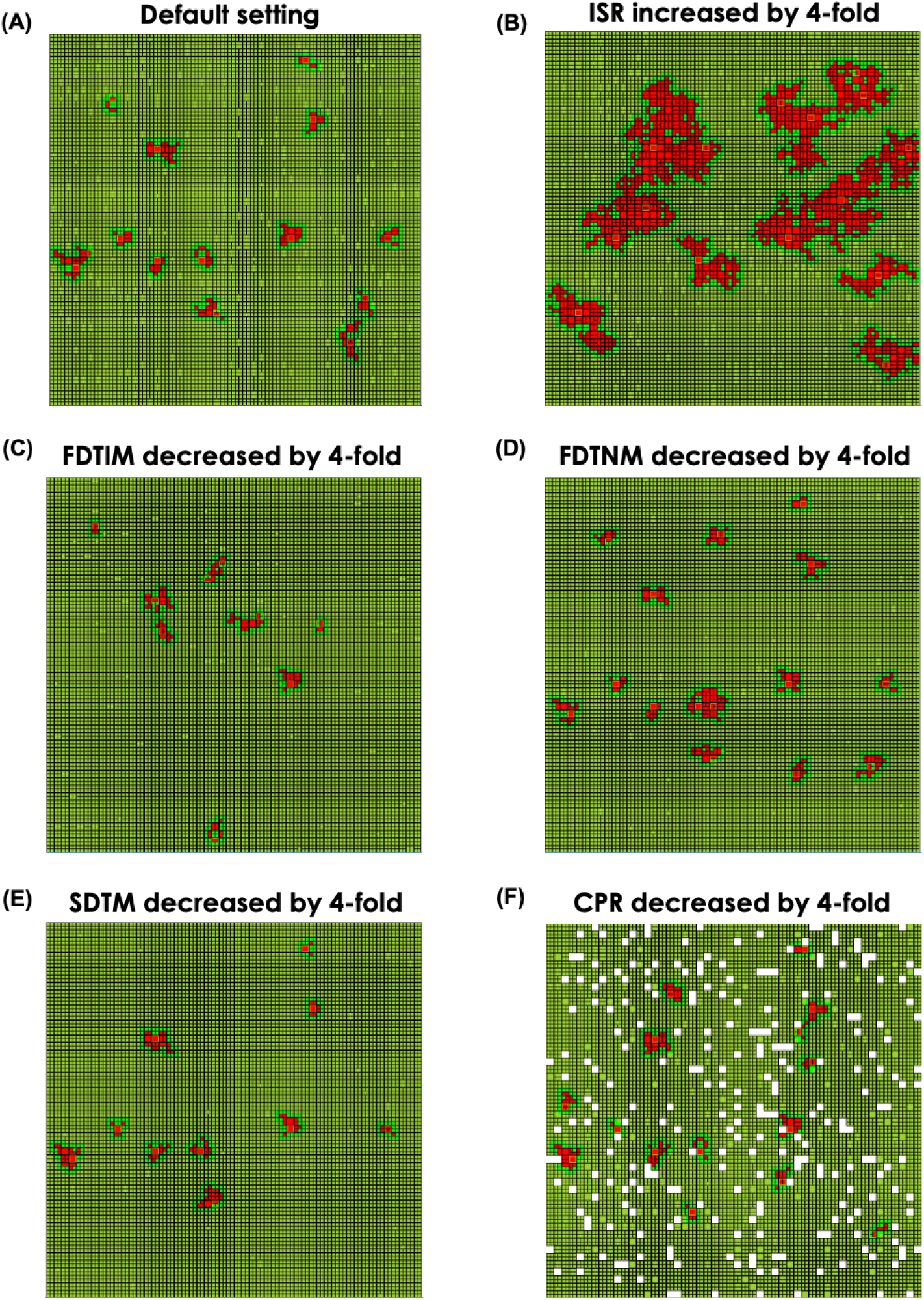
Effect of cell-to-cell infection spread rate and cell division on foci expansion. Image of foci formed *in silico* when (A) default cell division conditions are used (averaged foci size:10.3 ± 5.8); **(B)** infection spread rate (ISR) is increased (averaged foci size:181.9 ± 142.6); or when cell division is increased by **(C)** decreasing the time of the first division of initial cell (FDTM) (averaged foci size:10.0 ± 5.8); **(D)** decreasing the time of the first division of non-initial cell (FDTNM) (averaged foci size:12.9 ± 6.8); **(E)** decreasing the time of the second division (SDTM), or **(F)** decreasing cell proliferation to empty space (CPR) (averaged foci size:8.4 ± 5.0).

**FIG S2.**
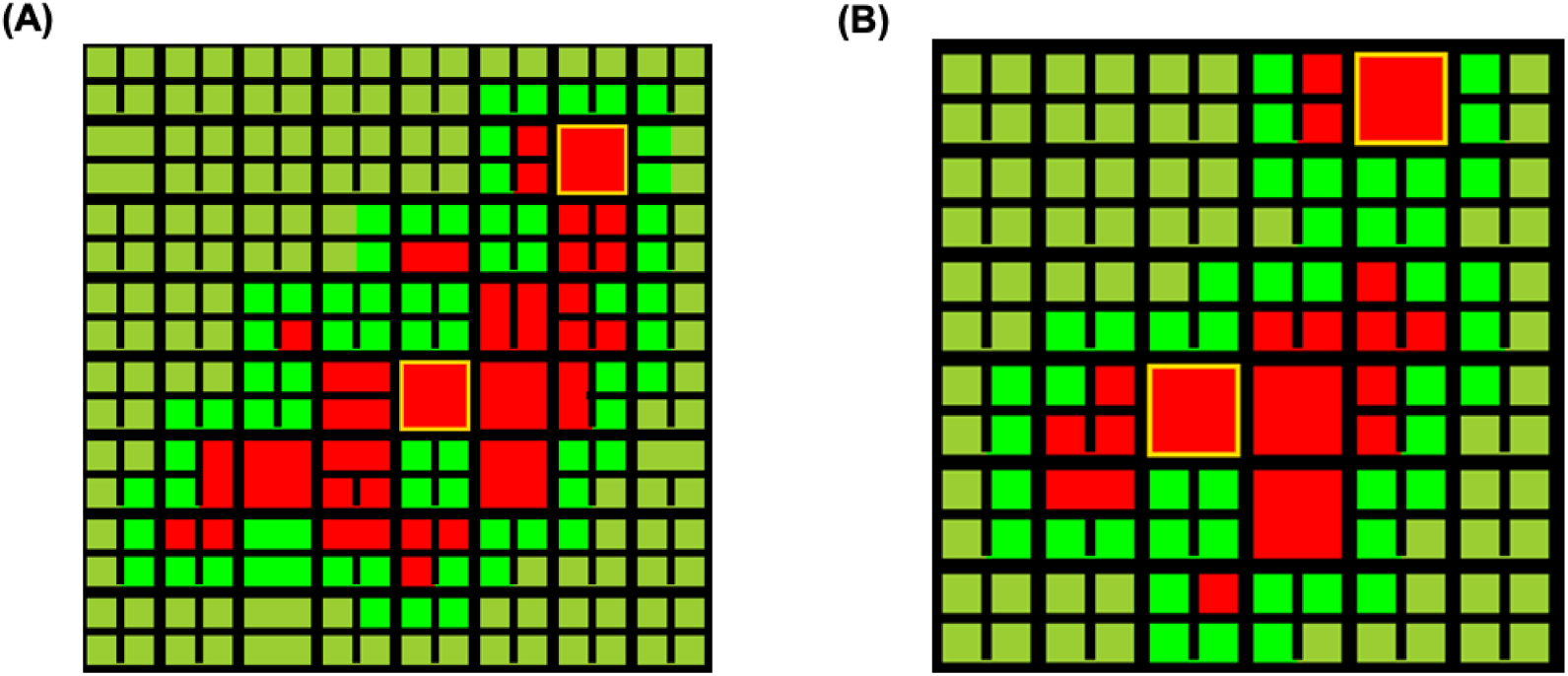
Effect of cell-division on foci merging. Image of merging foci with all variables set the same except for the second division (SDTM). Common parameters include: ISR = 10.86, CPR = 12.96, FDTIM = 34.31, FDTNM = 23.20. **(A)** SDTM =28.04. **(B)** SDTM = 2.04, which represents a 14-fold increase in rate of the second cell division.

**FIG S3.**
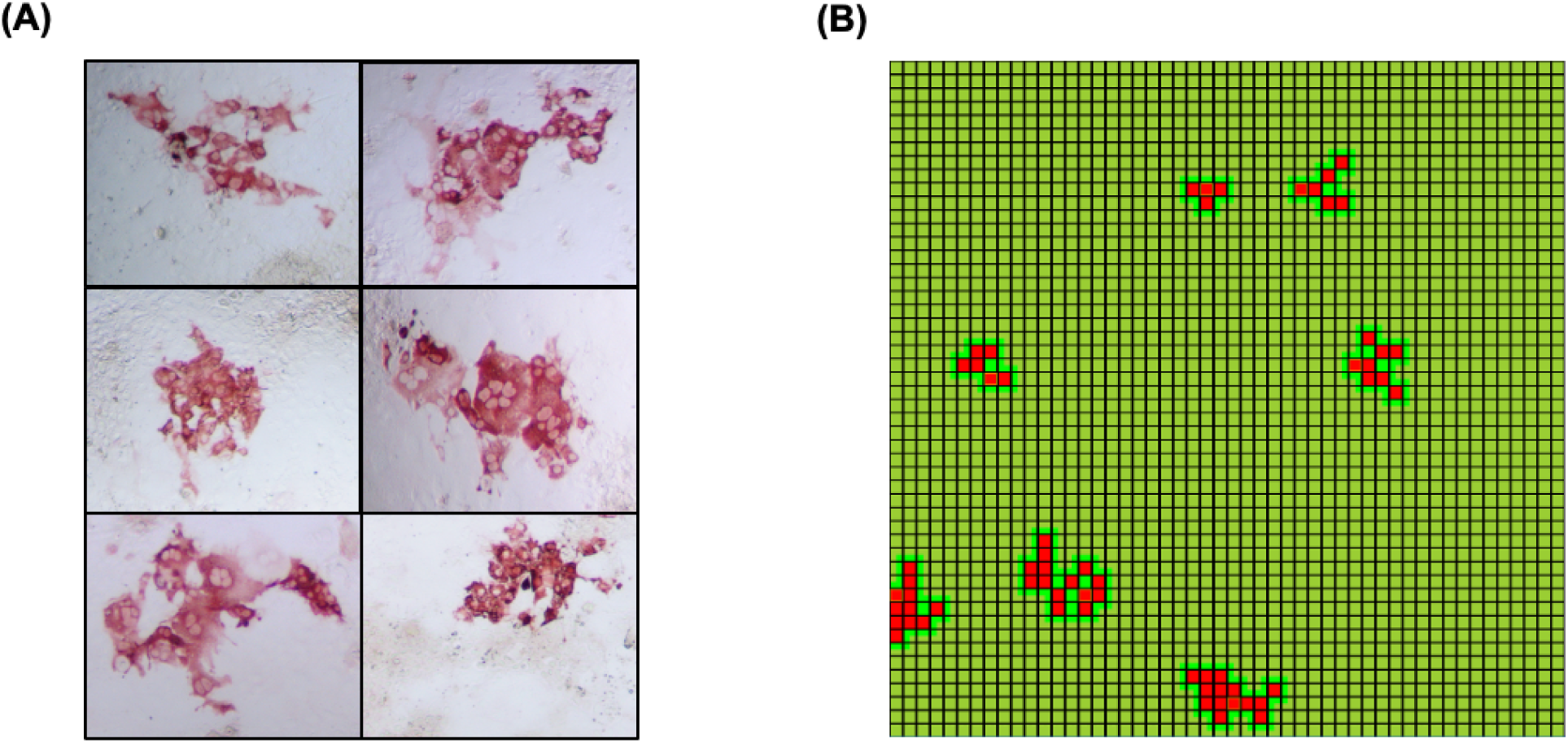
Irregular foci shape is observed in the absence of cell division. To determine if concurrent cell division and viral spread is responsible for the irregular HCV foci shape observed, infection was performed in non-dividing cells *in vitro* and *in silico*. **(A)** Foci in non-dividing cells in vitro. Huh7 cells were plated at confluence and incubated with 1% DMSO for 20 days to achieve a non-dividing state (7) before being infected for a CTC spread assay as described in Figure 1 (15). Infected cells were fixed and stained for HCV E2. **(B)** Foci in non-dividing cells in silico. Simulations were run using the parameter combination (ISR: 10.86; CPR: 12.96; FDTIM: 343100; FDTNM: 232000; SDTM: 280400), in which FDTIM, FDTNM, and SDTM was increased by 10000 times compared to the default parameter values in Figure 7 to achieve a non-dividing cell scenario prior to infection. Examples of simulated foci are shown.

